# Contrasting model mechanisms of alanine aminotransferase (ALT) release from damaged and necrotic hepatocytes as an example of general biomarker mechanisms

**DOI:** 10.1101/2019.12.24.887869

**Authors:** Andrew K. Smith, Glen E.P. Ropella, Mitchell R. McGill, Preethi Krishnan, Lopamudra Dutta, Ryan C. Kennedy, Hartmut Jaeschke, C. Anthony Hunt

## Abstract

Interpretations of elevated blood levels of alanine aminotransferase (ALT) for drug-induced liver injury often assume that the biomarker is released passively from dying cells. However, the mechanisms driving that release have not been explored experimentally. The usefulness of ALT and related biomarkers will improve by developing mechanism-based explanations of elevated levels that can be expanded and elaborated incrementally. We provide the means to challenge the ability of closely related concretized model mechanisms to generate patterns of simulated hepatic injury and ALT release that scale (or not) to be quantitatively similar to the wet-lab validation targets. The validation targets for this work are elevated measures of plasma ALT following acetaminophen (APAP) exposure in mice. We build on a published model mechanism that helps explain the generation of characteristic spatiotemporal features of APAP hepatotoxicity within hepatic lobules. Discrete event and agent-oriented software methods are most prominent. We instantiate and leverage a small constellation of concrete model mechanisms. Their details during execution help bring into focus ways in which particular sources of uncertainty become entangled within and across several levels with cause-effect details. Monte Carlo sampling and simulations comprise a virtual experiment. Falsification of one (or more) of the model mechanisms provides new knowledge and shrinks the model mechanism constellation incrementally. We challenge the sufficiency of four potentially explanatory theories for ALT release. The first model mechanism tested failed to achieve the initial validation target, but each of the three others succeeded. We scale ALT amounts in virtual mice directly to target plasma ALT measures in individual mice. Results for one of the three model mechanisms matched all target ALT measures quantitatively. We assert that the actual mechanisms responsible for ALT measures in individual mice and the virtual causal processes occurring during model execution are strongly analogous within and among real hepatic lobular levels.

**Author summary:** Interpretations of elevated biomarkers for drug-induced liver injury assume passive release during hepatocyte death, yet indirect evidence indicates that plasma levels can increase absent injury. Limitations on measurements make it infeasible to resolve causal linkages between drug disposition and plasma levels of biomarkers. To improve explanatory knowledge, we instantiate within virtual mice, plausible mechanism-based causal linkages between acetaminophen disposition and alanine aminotransferase (ALT) behavior that enables simulation results to meet stringent quantitative validation prerequisites. We challenge the sufficiency of four model mechanisms by scaling ALT measurements in virtual mice to corresponding plasma values. Virtual experiment results in which ALT release is a combined consequence of lobular-location-dependent hepatocyte death and drug-induced cellular damage, matches all validation targets. We assert that the actual mechanisms responsible for plasma ALT measures in individual mice and the virtual causal processes occurring during model execution are strongly analogous within and among real hepatic lobular levels.

## Introduction

The use of several conventional clinical biomarkers (e.g., alanine aminotransferase (ALT), aspartate aminotransferase, creatine kinase, lipase, cardiac troponin, etc.) assumes that the analytes are passively released from dying cells [1, 2]. However, the mechanisms have not been resolved experimentally. We seek improved understanding of the temporal events leading to and driving biomarker release to aid their interpretation, especially in cases where biomarker measures may be linked to medications, and to inform the identification, validation, and use of future biomarkers. We begin by focusing on ALT release from parenchymal cells (hepatocytes) in the liver. ALT is a good candidate for developing plausible release mechanisms. It has been measured in circulation since the 1950s [3] and has become the standard clinical biomarker of liver injury. Because transient ALT elevations are frequently observed during preclinical testing and clinical trials of new drugs, the correct interpretation of its measures can be critical [2]. There may be considerable diversity in the processes linking ALT release with different initiators of tissue damage. To enable challenging competing ALT release hypotheses, we start by focusing on acetaminophen (APAP), a widely used, well-studied analgesic that, when overdosed, causes hepatotoxicity. It is generally assumed that elevated measures of serum ALT are a direct consequence of release from hepatocytes undergoing necrosis, yet some patients treated with therapeutic doses of APAP can experience transient elevations absent any evidence of liver injury [4]. Large variations in serum ALT measures across preclinical models are another barrier to interpretation. For example, Harrill et al. [5] reported a 9- to 20-fold variation in mean serum ALT among different mouse strains that received a standard toxic APAP dose.

The main objective of this work is to develop, support, and challenge plausible cause-effect linkages between APAP disposition and metabolism and concurrent measurements of ALT in plasma. Doing so is a requisite for resolving plausible explanations for variation within and among studies and thereby diminishing barriers to interpretation. However, it is currently infeasible to establish such linkages in vivo because hepatocyte damage and resulting ALT release cannot be measured directly. Despite these barriers and uncertainties, progress is being made using conventional mathematical modeling methods. For example, Howell et al. [6] used a multifaceted drug-induced liver injury model to offer new insights into serum ALT measures from several studies. Their hypothetical ALT release sub-model assumes that necrotic cells release ALT. It describes released ALT moving to intralobular spaces and then into the blood.

Multi-source uncertainties have proven to be a stubborn barrier to progress in explaining ALT release. The underlying novel advantage provided by the approach and methods that we employ is the ability to represent both knowledge and ignorance concurrently by employing multiple computational methods. Discrete event and agent-oriented software methods are most prominent. We instantiate in software and leverage a small constellation of concrete model mechanisms that we intend to be strongly analogous to actual in vivo counterparts. We use the model mechanisms illustrated in Fig. 1 A-F as the parent mechanism because it has demonstrated scientific utility and quantitative biomimicry by achieving prespecified validation targets for particular use cases [7]. Unlike the conventional computational modeling methods used by Howell et al. [6] and others, making predictions is not our primary objective. Instead, we provide the means to challenge the ability of closely related model mechanisms to generate patterns of hepatic injury and ALT release that scale (or not) to be quantitatively similar to validation targets.

**Fig. 1.**
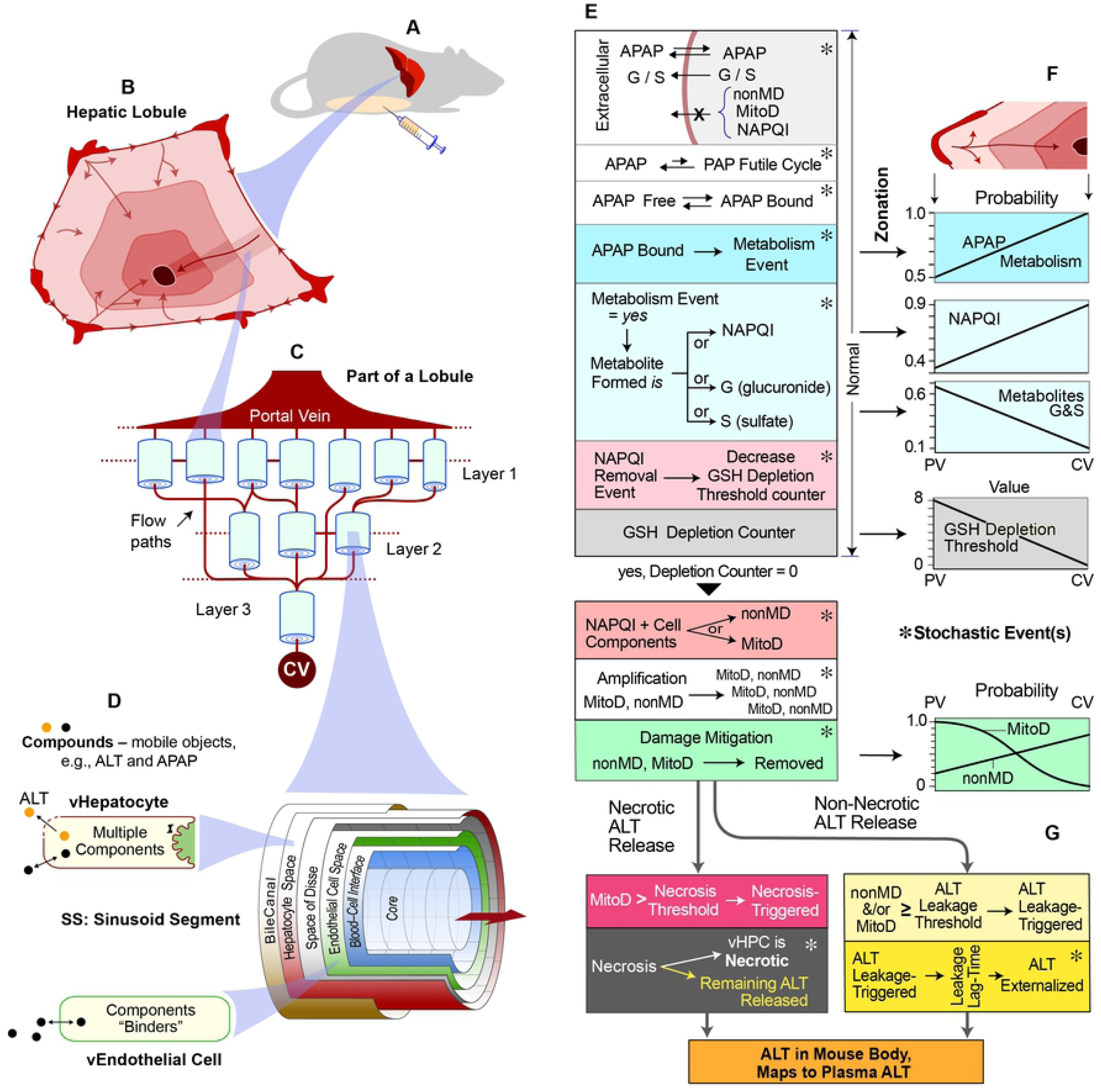
Component organization and model mechanism features. (**A**) A virtual Mouse, detailed in *Methods*, is a concretized, coarse-grain software analogy of an actual mouse. (**B**) Shading illustrates idealized periportal (PP) to pericentral (PC) gradients within a cross-section of a hepatic lobule. (**C**) A portion (16%) of one virtual Lobule is illustrated. A Monte-Carlo specified interconnected directed graph, which can be different (within constraints) for each experiment, specifies flow paths for APAP and other Solutes (see Methods). (**D**) A multi-layered Sinusoid Segment (SS) maps to a portion of hepatic tissue. One is placed at each graph node. A space within an SS contain Virtual Hepatocytes (vHPCs), which contain a variety of objects needed to enable the cause-effect events within the model mechanism. (**E**) Each of these events and activities (along with those in **F** and **G**) may occur each simulation cycle. Events in the top series, labeled “Normal,” occur following therapeutic (non-toxic) and toxic doses of APAP. Those listed subsequently contribute to simulated APAP-induced injury. (**F**) Parameterizations of these features have the indicated Lobule-location dependency and are the same as in Smith et al. (7). There is a direct mapping between the probability of an APAP Metabolism event and average metabolic capacity at various lobular locations. Each vHPC uses location-dependent values drawn from those gradients. A No-ALT-Model a virtual Mouse using only parameterized variant of **A**-**F**. (**G**) The logic of the two ALT release processes is shown. The value for ALT Leakage Threshold is independent of Lobular location. Once ALT Leakage is Triggered, a value for ALT Leakage Delay is determined. The minimum ALT Leakage Lag-Time is < than the minimum Necrosis interval. Externalized ALT objects exit a SS, enter Central Vein (CV), and are transferred to Mouse Body the same as unmetabolized APAP. The mouse counterparts to these events and features are independent of APAP because the mouse counterparts of nonMD and MitoD can be generated in many ways.

Model mechanism details during execution help bring into focus ways in which particular sources of uncertainty become entangled with cause-effect details. Repetitive executions comprise an experiment, which challenges the hypothesis that a particular model mechanism will achieve prespecified target phenomena. Observing the unfolding of events during execution allows one to think through the model mechanism’s networked cause-effect details and identify (or not) weaknesses. Falsification of one (or more) of the model mechanisms—demonstrating its inability to achieve its target phenomena—provides new knowledge and shrinks the constellation of explanatory model mechanisms incrementally.

The target phenomena for this work are elevated plasma ALT measures following APAP exposure in mice [8]. We challenge the sufficiency of four potentially explanatory theories using virtual experiment methods [7, 9, 10]. The first model mechanism limits ALT release to necrotic cells. It failed to achieve the initial validation target, but each of the three other mechanisms succeeded. These included a second, necrosis-independent ALT release process that is triggered by lobular-location-dependent APAP-induced damage products. We scale ALT in virtual mice directly to individual target plasma ALT measures. Results for one of the three model mechanisms matched all target ALT measures quantitatively. It is the first cause-effect model mechanism to provide plausible explanations for entanglements of APAP metabolism and hepatic disposition, accumulation of toxic damage, externalization of ALT from hepatocytes, and ALT accumulation in plasma following a toxic APAP dose in mice.

As the knowledge instantiated within these model mechanisms increases, they can be leveraged to improve and individualize the utility of biomarkers of drug-induced liver injury and to understand the sources of interindividual variability better. We assert that the actual mechanisms responsible for the individual plasma ALT measures during the targeted study and the virtual causal processes occurring during model mechanism executions are dynamically analogous within and among lobular levels. The modularity of virtual mouse components and processes, from mobile APAP objects to metabolism and formation of damage products within individual hepatocytes, to virtual liver architecture, to mouse body, is designed to enable seamless translation to humans and extend use cases to other toxicants and biomarkers.

## Results

### The same parent Hepatotoxicity Mechanism for all experiments

A virtual experiment is a test (a trial) of an extant model mechanism (MM) hypothesis (see *Model mechanism requirements*). To distinguish virtual mouse components, characteristics, and phenomena from real counterparts, we capitalize the former hereafter and, in three cases, append the prefix “v”. Names of parameters are italicized.

We used a parameterized version of the MM illustrated in Fig. 1A-1F to explain how and why periportal (PP) to pericentral (PC) gradients of three processes is necessary and sufficient to explain the characteristic PC occurrence of necrosis following a toxic APAP dose in mice [7]: 1) production of the reactive metabolite N-acetyl-p-benzoquinone imine (NAPQI), 2) glutathione (GSH) depletion, and 3) mitigation of mitochondrial damage. Results of virtual experiments support the claim that, at corresponding degrees of granularity, the MM during execution and the actual APAP-induced hepatotoxicity mechanism may be strongly analogous within and across multiple lobular levels. That evidence supports using that MM as the parent MM for this work. For the results that follow, parameterizations of components in Fig. 1A-1D, and events in Fig. 1E and 1F are unchanged from Smith et al. [7]. Parameterizations for all key features are listed in Supporting S1 Table. Achieving the validation targets for this work without having to alter the parent MM will strengthen the case that it is strongly analogous to the actual APAP-induced hepatotoxicity mechanism. However, failure to achieve those validation targets risks falsifying the parent MM.

### Two ALT release processes: one confirmed, the other plausible

Evidence suggests externalization of damaged macromolecules is a normal, ongoing hepatocyte process. Additional externalization processes may (or not) be engaged as rates of damage product accumulation increase and the nature of those products change. Consequently, undamaged macromolecules such as ALT may become coupled to one or more of these processes and become externalized directly or concomitantly.

The parsimonious working hypothesis, illustrated in Fig. 2A, is that ALT may be released by two separate processes, passive release as a consequence of necrosis and non-necrotic release processes. The supporting evidence is circumstantial. Gamal et al. [11] described leakage of cytoplasmic material associated with APAP-induced disruption of hepatocyte integrity and surface damage associated with blebbing. Ni et al. [12] reported that hepatocytes upstream of necrosis experience hepatocellular damage mitigation, including evidence of autophagy and mitophagy. Following a subtoxic APAP dose, APAP-protein adducts are externalized to blood in the absence of necrosis [8, 13]. If a normal cellular process is damaged during APAP-induced injury, its selectivity may be eroded leading to concomitant externalization of other macromolecules, including ALT. Consistent with that scenario, McGill et al. [8] reported that, following a low toxic APAP dose, APAP-protein adducts are externalized to blood prior to ALT elevations.

**Fig. 2.**
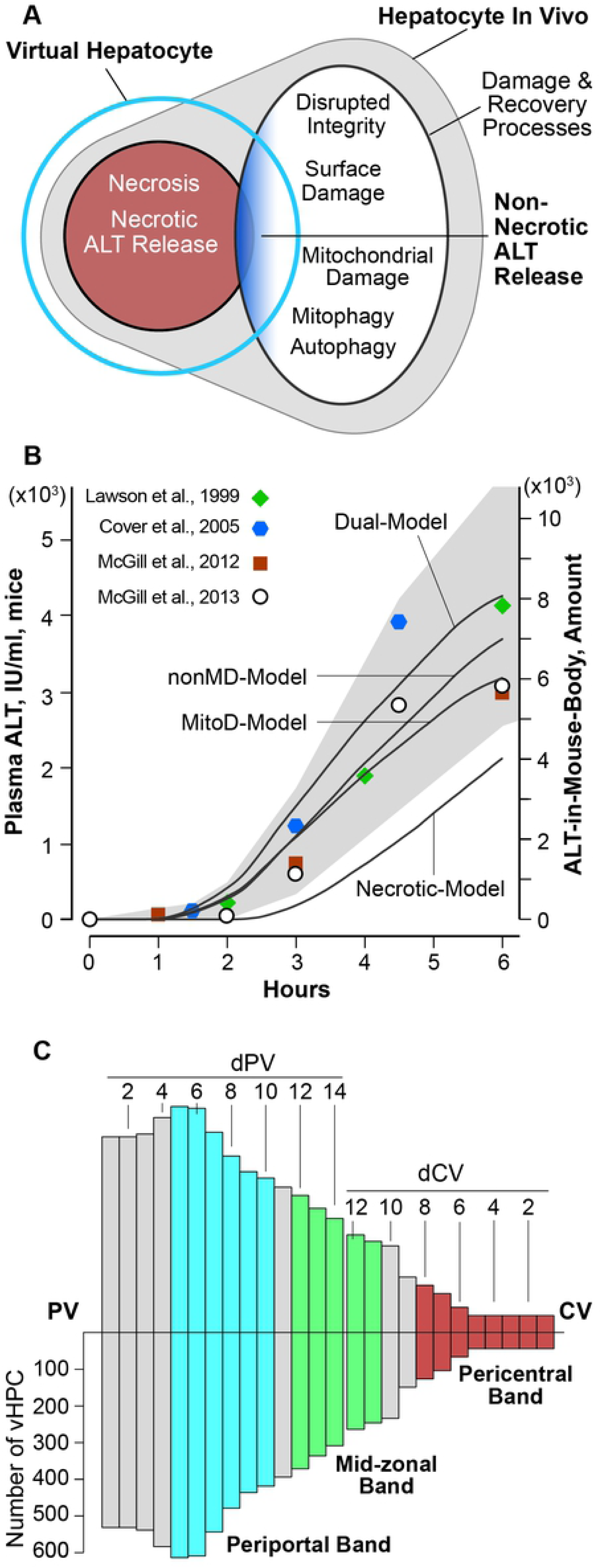
ALT release processes, validation targets, and Lobular organization. (**A**) An illustration of plausible relationships between two virtual ALT release processes and in vivo counterparts. Necrotic ALT Release (Fig. 1G) maps to confirmed ALT release during necrosis. Non-Necrotic ALT Release maps to a structured conflation of non-necrotic damage and recovery processes that may directly or concomitantly enable ALT release, indicated by blue shading. (**B**) The gray area is the initial validation target range (left axis), which is based on mouse data from the four indicated reports. The minimal Similarity Criterion for an acceptable ALT release model mechanism (MM) is that it generates ALT-in-Mouse-Body profiles that, when scaled (right axis), fall within the target range. The four labeled profiles were generated by the MMs described in the text. (**C**) Bar heights represent the mean number of vHPCs at the indicated location within an average vLobule. The left edge corresponds to Portal Vein (PV). The right edge corresponds CV. Moving left-to-right, the first 14 bars are located at increasing distances from PV (dPV) along the average PV-to-CV path. Moving right-to-left, the first 12 bars are located at increasing distances from CV (dCV) along the average CV-to-PV path. PP band = blue bars, Mid-Zonal (M-Z) band = green bars, and PC = red bars.

We conjectured that instantiating a virtual counterpart to non-necrotic ALT release processes (see *Representing ALT release from damaged hepatocytes*) should be feasible without requiring any change to the parent MM because the two types of damage products generated (MitoD and nonMD) map to a coarse grain conflation of all types of hepatocellular damage that could lead to ALT release [7].

### Necrosis-Model: simulating ALT release from hepatocytes undergoing necrosis

The parent MM spanning Fig. 1A-1F (the No-ALT-Model) includes transitioning a virtual hepatocyte (vHPC) from the Necrosis-Triggered state to the Necrotic state following a Necrosis interval. Instantiating a virtual counterpart to ALT release caused by necrosis was straightforward. We added ALT objects to each vHPC along with the instruction that any remaining ALT objects are released when the vHPC becomes Necrotic. Because there is no strong wet-lab evidence to the contrary, we assumed that each hepatocyte, independent of lobular location, contains the same amount of ALT. Using ALT per vHPC = 5 objects at time (t) = 0 proved sufficient for this work. One ALT object maps to a small amount of actual ALT. When a vHPC transitions from Necrosis-Triggered to Necrotic, all remaining ALT is released and externalized (bottom, Fig. 1G). Externalized ALT objects follow the same stochastic movement rules as APAP, G (the virtual counterpart to the glucuronide metabolite of APAP), and S (the virtual counterpart to the sulfate metabolite) [7]. Upon entering the Central Vein (CV), an ALT is moved to Mouse Body. An ALT in Mouse-Body is scaled directly to represent plasma ALT. We name that entire sequence, from Metabolism to ALT externalization, the Necrosis-Model. Following a tissue distribution phase, the plasma half-life for ALT in mice is approximately 25 h [14]. During the first 6 h of experiments in mice, we conjecture that little of the externalized ALT is cleared. For simplicity, ALT in Mouse-Body is not removed. It accumulates and does not re-enter the vLiver.

### Explaining ALT release from damaged hepatocytes using three model mechanism variants

Early workflow results suggested that ALT externalization and accumulation in Mouse Body as a consequence of Necrosis-Model executions could not scale quantitatively to closely match the temporal patterns of plasma ALT, the validation targets. Accordingly, we sought parsimonious extensions of the parent MM that 1) could map to the non-necrotic ALT release process, 2) would operate concurrently with—but independent of—Necrosis-Model, and 3) make ALT Externalization a direct function of Damage Products. The parent MM in Fig. 1E includes generation of two types of Damage Products, nonMD (maps to all types of non-mitochondrial damage products) and MitoD (maps to all types of mitochondria related damage products) within each vHPC along with their concurrent removal via Damage Mitigation. Those events are independent of Necrosis-Triggered. Together, they map to the Damage and Recovery Processes in Fig. 2A. We instantiated three versions of the coarse grain Non-Necrotic ALT Release process illustrated in Fig. 1G by making ALT externalization a direct function of amount of Damage Products in each vHPC: nonMD, MitoD, or both. When MitoD (nonMD) is the cause of ALT Externalization, we name the MM MitoD-Model (nonMD-Model). When both cause of ALT Externalization, we name it the Dual-Model.

Each simulation cycle, each vHPC determines if its amount of designated Damage Products (nonMD, MitoD, or both) exceeds an ALT Leakage Threshold value. If so, that vHPC becomes Leakage-Triggered and initiates an ALT externalization process. In MitoD-Model (nonMD-Model), only the amount of MitoD (nonMD) within each vHPC is compared to the Leakage Threshold value. In Dual-Model, the sum of nonMD and MitoD is compared to the Leakage Threshold each simulation cycle. Results from experiments to achieve stringent validation targets (described below) used MitoD-Model because we judged it to be the more parsimonious of the three. We also observed no explanatory advantage when using nonMD-Model or Dual-Model.

The vHPC specifies a Leakage Lag-Time by a pseudo-random draw from a uniform [Min, Max) distribution (see *Kinetics of ALT externalization*). When the Lag-Time duration is reached and the vHPC’s ALT counter value > 0, the vHPC creates an ALT object, externalizes it, and decrements its ALT counter by 1. An externalized ALT in “Blood” exits the CV by following the same movement rules used by APAP Metabolites.

The initial value of the ALT counter is 5. As stated above, using ALT objects per vHPC = 5 proved sufficient for this work. We could have added five ALT objects to each vHPC at t = 0. Although not biomimetic, creating ALT objects as they are needed is computationally more efficient. Tracking objects in all vHPCs that are doing nothing is computationally inefficient and increases the duration of executions. Once an ALT externalization is scheduled, it will occur at that time if the ALT counter value > 0. After an ALT externalization is scheduled, the designated Damage Products may fall below the Leakage Threshold. When that occurs, the vHPC is no longer Leakage-Triggered.

We conducted exploratory experiments using Leakage Threshold values ≤ 5. Setting the Threshold = 1, independent of Portal Vein (PV) to CV location, enabled achieving validation targets. For Necrosis-Model, when a Necrosis-Triggered vHPC transitions to Necrotic, and its ALT counter value > 0, the vHPC creates ALT objects corresponding to the counter value and externalizes them with zero-time delay.

### Specifying the initial validation target range

Although the above four MM variants are coarse grain analogies of the referent processes, the constellation of feature and event parameterizations that might merit exploration is large, and there are too few wet-lab data to guide selecting particular parameter value combinations. Thus, an essential early workflow goal was to identify a subset of that constellation on which to focus (see “*Conducting many narrowly focused virtual experiments*”). To do so, we specified a broad, semiquantitative validation target: the temporal trend of ALT-in-Mouse-Body values must exhibit a sigmoidal shape and fall within the target area in Fig. 2B. Specification of the upper and lower bounds of that target region was guided by a selection of mean plasma ALT measures from four mouse studies that studied toxic APAP doses [8, 15–17]. Two studies used C3Heb/FeJ mice [15, 16] and two used C57BL/6 mice [8, 17]. Three studies used a 300 mg/kg APAP dose, whereas one used a 500 mg/kg dose [15]. Results from the 500 mg/kg APAP studies were included because there is considerable variance among mouse strains in sensitivity to APAP-induced hepatotoxicity, and we hypothesize that, when responding to a comparably toxic APAP dose, a common set of entangled mechanism features governs ALT release and its subsequent accumulation in plasma.

To acknowledge the variability within and between studies, we specified that the upper edge of the target region shall include the mean plus the value of 1 standard-error-of-the-mean for the 3-to-6 plasma ALT measures. We also specified that the lower edge of the target region shall include the mean minus the value of 1 standard-error-of-the-mean of those values.

### Intralobular location-dependent measurements

Because the temporal pattern of upstream events can significantly influence the temporal pattern of downstream events, we record temporal measurements of selected features within the three bands illustrated in Fig. 2C.

During each execution, there is a large variety of PV-to-CV paths that an APAP object might travel. The distribution of mean PV-to-CV distances is significantly skewed toward longer distances to validate against single-pass liver perfusion data for multiple drugs [18]. We use the length of the average upstream lobular Layer (in Core grid points) to specify the distance of each vHPC from PV (designated dPV). Averaging over the Monte Carlo executions within an experiment, we determine the mean number of vHPCs at each dPV location. However, because we are particularly interested in the temporal order of events occurring within pericentral (PC) vHPC, we also identify vHPC locations in terms of distance from CV. The distribution of CV-to-PV distances is different from the PV-to-CV distribution because the Sinusoidal Segments (SSs) in Fig. 1A in each layer can have different lengths and one upstream SS can link to multiple downstream SSs. Nevertheless, we specify the location of each vHPC along the average SS path as a distance from CV (designated dCV). At each dCV location we determine the mean number of vHPCs. Figure 2C combines a visualization of the mean number of vHPCs at dPV locations 1 through 14, which accounts for 85.27% of the average number of vHPCs per experiment, with mean number of vHPCs at dCV locations 1-12, which accounts for 14.14% of the average number of vHPCs. The combined visualization accounts for 99.41% of average total vHPCs. On average, for the detailed result described below for MitoD-Model, the PP band includes 4772 vHPCs that are 5-10 grid points from PV (blue bars in Fig. 2C). The Mid-Zonal (M-Z) band includes 1721 vHPCs that are 12-16 grid points from PV (green), and the PC band includes 906 vHPCs that are 0-8 grid points from CV (red bars).

During each execution, for each vHPC, we measure amounts of APAP, G, S, NAPQI, nonMD, and MitoD, along with counts of the following events: GSH Depletion, Damage Mitigation, Leakage-Triggered (defined below), ALT externalized (described below), Necrosis-Triggered, and Necrotic. Those values are summed or averaged over all Monte Carlo (MC) executions and within the three bands illustrated in Fig. 2C.

### APAP disposition and toxicity measurements

APAP disposition and toxicity measurements for MitoD-Model are provided in Fig. 3. Corresponding measurements for Necrosis-Model, Dual-Model, and nonMD-Model are identical to those for the MitoD-Model within the variability of Monte Carlo executions. The measurements in Fig. 3B show that the amount of APAP per vHPC increases PP-to-PC. That increase is a direct consequence of fewer PC vHPCs (Fig. 2C) being exposed to the amount of incoming APAP. NAPQI differences among the three bands (Fig. 3C) are more dramatic than those for APAP because the fraction of APAP that is Metabolized to NAPQI, rather than to G and S Metabolites, increases PP-to-PC (Fig. 1F). Figure 3D shows the average location of Necrosis-Triggered events within PC and M-Z bands, independent of the number of such events (Fig. 3I). The order of the cumulative GSH Depletion profiles in Fig. 3E may seem inconsistent with the order of amounts in Fig. 3, B and C. The explanation is that by 1 h post-Dose, the GSH Depletion Threshold for a majority of vHPCs within the PC band has been breached. However, GSH Depletion Thresholds within the M-Z band are larger (Fig. 1F), so GSH Depletion continues even though less NAPQI per APAP is being formed.

**Fig. 3.**
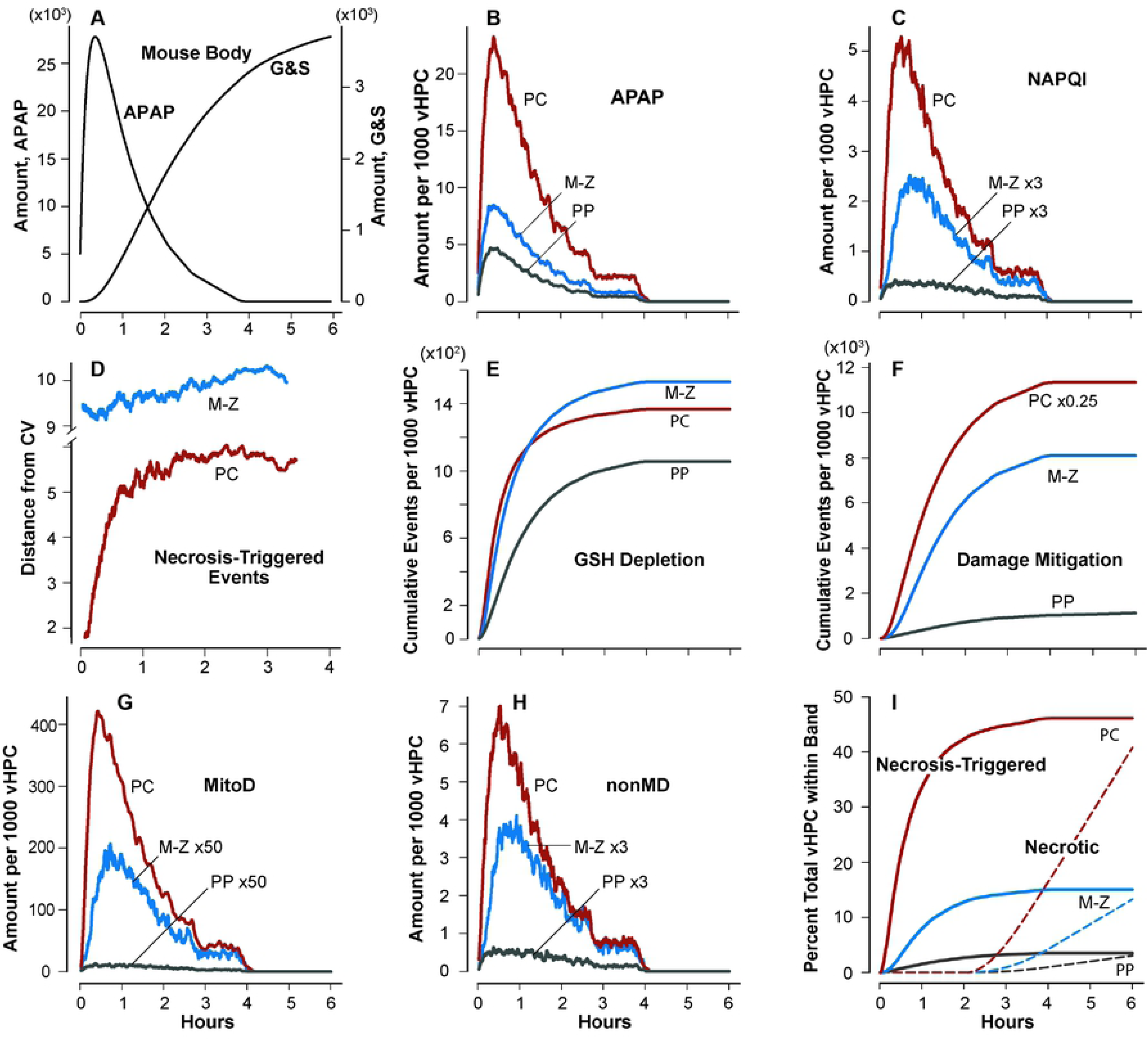
Temporal profiles of measurements made during executions of the parent model mechanism. (**A**) APAP and its Metabolites, G and S, in Mouse Body. Values in **B**-**E** are centered moving averages (181-point). (**B**) Average amount of APAP per 1000 vHPCs within the PP, Mid-Zonal (M-Z), and PC bands identified in Fig. 2C. (**C**) Average amount of NAPQI within PP, M-Z, and PC bands. (**D**) Distance from CV of average Necrosis-Triggered events within PC and M-Z. The few events within the PP band are more distant. (**E**) Cumulative GSH depletion events. (**F**) Cumulative Damage Mitigation events. Average amount of MitoD (**G**) and nonMD (**H**) within PP, M-Z, and PC bands. (**I**) Cumulative Necrosis-Triggered and Necrotic events (dashed curves) within the three bands. All Necrosis-Triggered vHPCs become Necrotic following a Monte Carlo sampled Necrosis interval.

The amount of MitoD per vHPC within M-Z and PP bands (Fig. 3G) is minute compared to the amounts within the PC band. Three MM features, illustrated in Fig. 1E, help explain the differences. 1) The probability that an APAP Metabolism event will generate a NAPQI (reflected in Fig. 3, B and C) is smaller in M-Z and PP bands than in the PC band. 2) The GSH depletion threshold, which maps to relative amounts of GSH, decreases PP-to-PC. 3) A more influential feature is that the probability of a MitoD Damage Mitigation event (Fig. 3F) is larger in M-Z and PP bands than in the PC band. The amounts of nonMD within a vHPC in the PC band (Fig. 3H) are much smaller than MitoD amounts primarily because the probability of nonMD removal by Damage Mitigation is largest within the PC band, whereas, the probability of MitoD removal by Damage Mitigation is the smallest within the PC band (Fig. 1F). Within the PC band, 50% of Necrosis-Triggered events occur within the first 40 minutes (Fig. 3I). However, the minimum delay (Fig. 3I) before a Necrosis-Triggered vHPC becomes identifiable as Necrotic is 2 h.

### Quantitative mapping of ALT-in-Mouse-Body to individual plasma ALT measures

The most stringent validation targets are the plasma ALT measures from 18 matched mice that received the same toxic APAP dose during the same experiment [8]. To support a strong explanatory analogy, a MM must use the following direct mapping functions. At each measurement time, *t*, ALT objects in Mouse Body will scale directly to a fixed concentration (amount) of ALT in small aliquot of plasma (and vice versa). Specifically,

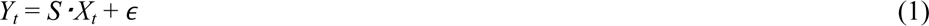

where *Y_t_* = the mean concentration of ALT in plasma (IU/ml) at time *t*; *X_t_* = the mean number of ALT objects in Mouse Body at *t* simulation cycles after Dosing; *S* is the scaling constant, and *ϵ* is random error. To account for individual variability, we hypothesize that a plasma ALT measurement at time *t* for an individual mouse is also directly proportional to the mean total ALT-in-Mouse-Body, as specified by

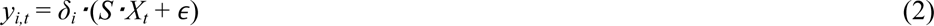

where *y_i,t_* = the measured concentration of ALT in plasma (IU/ml) for mouse *i* at time *t* after dosing, and *δ_i_* = the degree to which a mean ALT-in-Mouse Body value must be skewed (amplified or diminished) to match the individual’s plasma ALT value. *δ_i_* is applied to (*S·X_t_* + *ϵ*) because it seems infeasible to separate the individual random error.

The Individualized Mapping Criterion is most stringent. It specifies that *δ_i_* should be distributed somewhat symmetrically with mean = 1.0 (± tolerance) for all 18 mice. Absent comparable work to provide guidance, we started by setting tolerance arbitrarily at ± 0.1.

### Kinetics of ALT externalization

Elevated plasma (or serum) ALT measures are seen in humans and occasionally in mice following APAP doses considered to be nontoxic. From that, we inferred that the lag-time between formation of NAPQI reaction products (or NAPQI-induced damage products) and ALT externalization must be shorter than the lag-time between triggering and measuring necrosis. However, we lacked information to guide specification of the Min-Max range of the Leakage Lag-Time distribution.

Results of early explorations showed that ALT-in-Mouse-Body values are particularly sensitive to values chosen for Leakage Lag-Time. However, we had no information to guide specification of the Lag-Time distribution. Selection of a plausible minimum value was guided by whether or not scaled ALT-in-Mouse-Body profiles fell within the initial target range in Fig. 2B. We explored a variety of distribution ranges. By using the uniform distribution with [Min, Max) = [2700, 18000) simulation cycles, which scales to [0.75, 5] h (mean = 2.88 h), and a narrow range of Equation 1 *S* values, we were able to meet the initial target range criterion for MitoD-Model, nonMD-Model and Dual-Model, but not Necrosis-Model. We continued using that range for the duration of this work.

Although the disposition of APAP and the occurrence of Damage was identical among the four MMs, ALT release was different (Fig. 4A). The temporal pattern for accumulation of ALT-in-Mouse-Body from Necrosis-Model differed from those of three other MMs. Those differences result from differences in the rate of ALT exiting CV, which is a product of when and where ALT is externalized upstream of CV (Fig. 4B and 4C). Differences in amounts of ALT to be released within the three bands are provided in Fig. 4, D-F. By 30 min post-Dose, about 50% of vHPCs within the PC band have experienced a Leakage-Triggered event. Evidence of ALT externalization, which is controlled by Leakage Lag-Time, is seen a few minutes later (Fig. 4B). Entering Mouse Body is the last event. Consequently, any meaningful change within the networked casual process from Metabolism to Damage production and Damage Mitigation to ALT externalization can impact the shape of the resulting Body profile. Changes in Histological details, such as SS features (e.g. Cell spaces/densities, Space of Disse) along with dimensions and flow path connections, can also alter the ALT-in-Mouse-Body profile, absent any changes in events occurring within vHPCs.

**Fig. 4.**
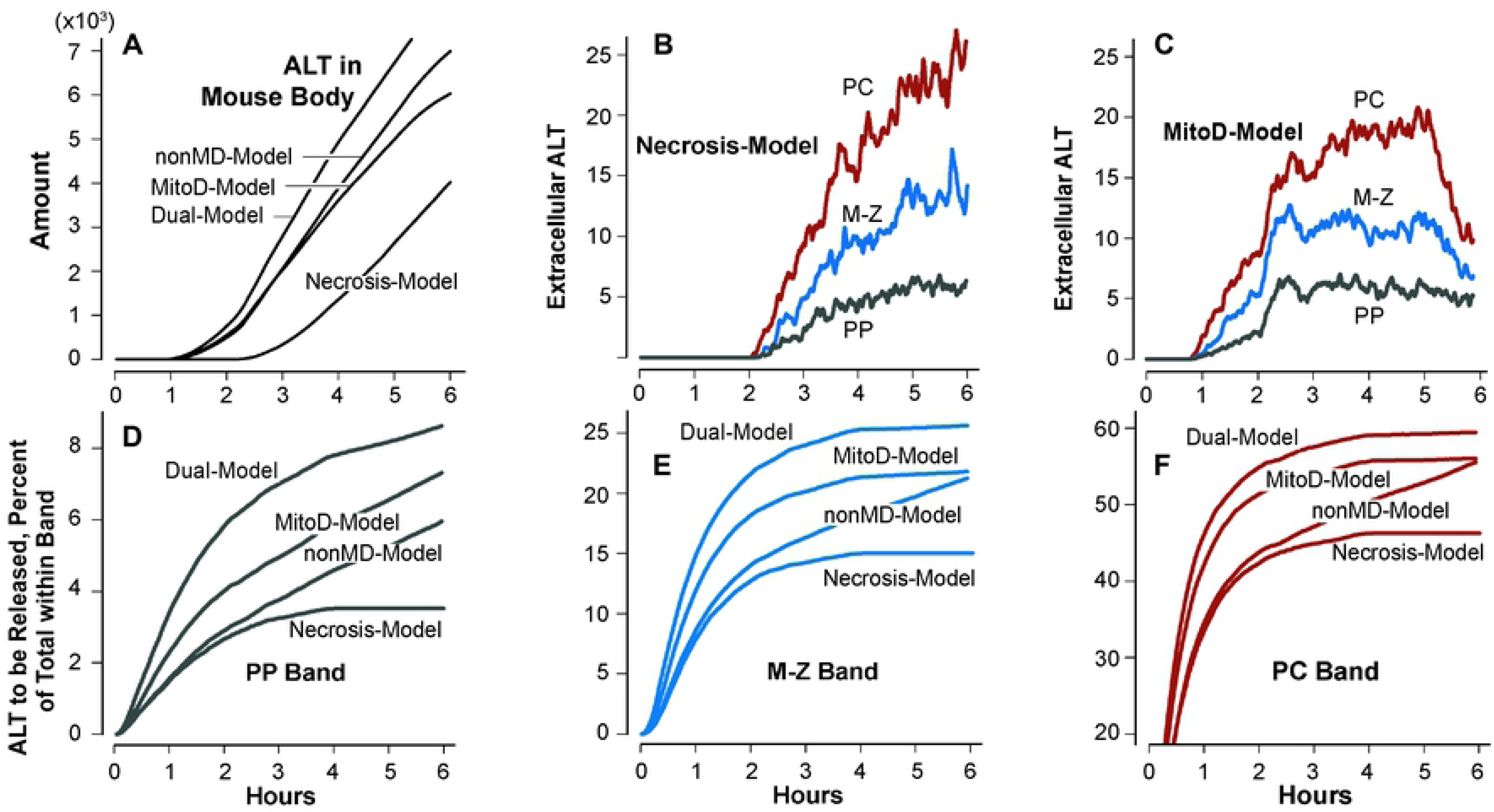
ALT Release characteristics and features during MM executions. (**A**) ALT accumulation in Mouse Body for each of the four MMs. (**B** and **C**) The measures are amount of ALT that has been externalized by vHPCs within the three Fig. 2C bands, but, at the time measured, has not yet exited the band during execution of Necrosis-Model (**B**) and MitoD-Model (**C**). The relative patterns of corresponding measures during execution of nonMD-Model and Dual-Model are similar to those for MitoD-Model. (**D**-**F**) Each panel contains data from one of the three bands. Within a panel, each profile is the cumulative percent of ALT that was scheduled for release at t or earlier. Once an ALT is scheduled for release, the event will occur following a Monte Carlo sampled Lag-Time.

### Individualized model mechanism-based explanation for plasma ALT measures

From an APAP hepatotoxicity perspective, we reasoned that MitoD-Model is marginally more parsimonious than nonMD-Model because MitoD production, rather than nonMD production, is directly correlated with observed PC patterns of tissue damage [7]. Further, MitoD-Model is more parsimonious than Dual-Model because ALT Leakage-Triggered within the latter is dependent on both categories of damage product. Accordingly, we used MitoD-Model to seek Eq. 2 parameterizations capable of achieving the Individualized Mapping Similarity Criterion. Should it be needed, the same method can be used to seek Eq. 2 parameterizations for nonMD-Model and Dual-Model that also achieve the Similarity Criterion.

At the start of the scaling workflow, we encountered an obstacle. The variability in plasma ALT measures at 4.5 and 6 h in Fig. 5A are consistent with commonly encountered interindividual variability in APAP-induced hepatotoxicity. However, the clustering of plasma ALT measures at 3 h (one subgroup of four mice; a second subgroup of two mice) suggests an additional influence that is absent from the mice measured at 4.5 and 6 h. We temporarily set aside the values for mice 3-6, and focused on the plasma ALT measures from the remaining 14 mice. We determined that using *S* = 1.72 achieved the Individualized Mapping Criterion: we obtained sample mean, *x̄* = 1.016, standard deviation, *s* = 0.277, and coefficient of variation = 0.274. The individualized mappings are shown in Fig. 5, B-D.

**Fig. 5.**
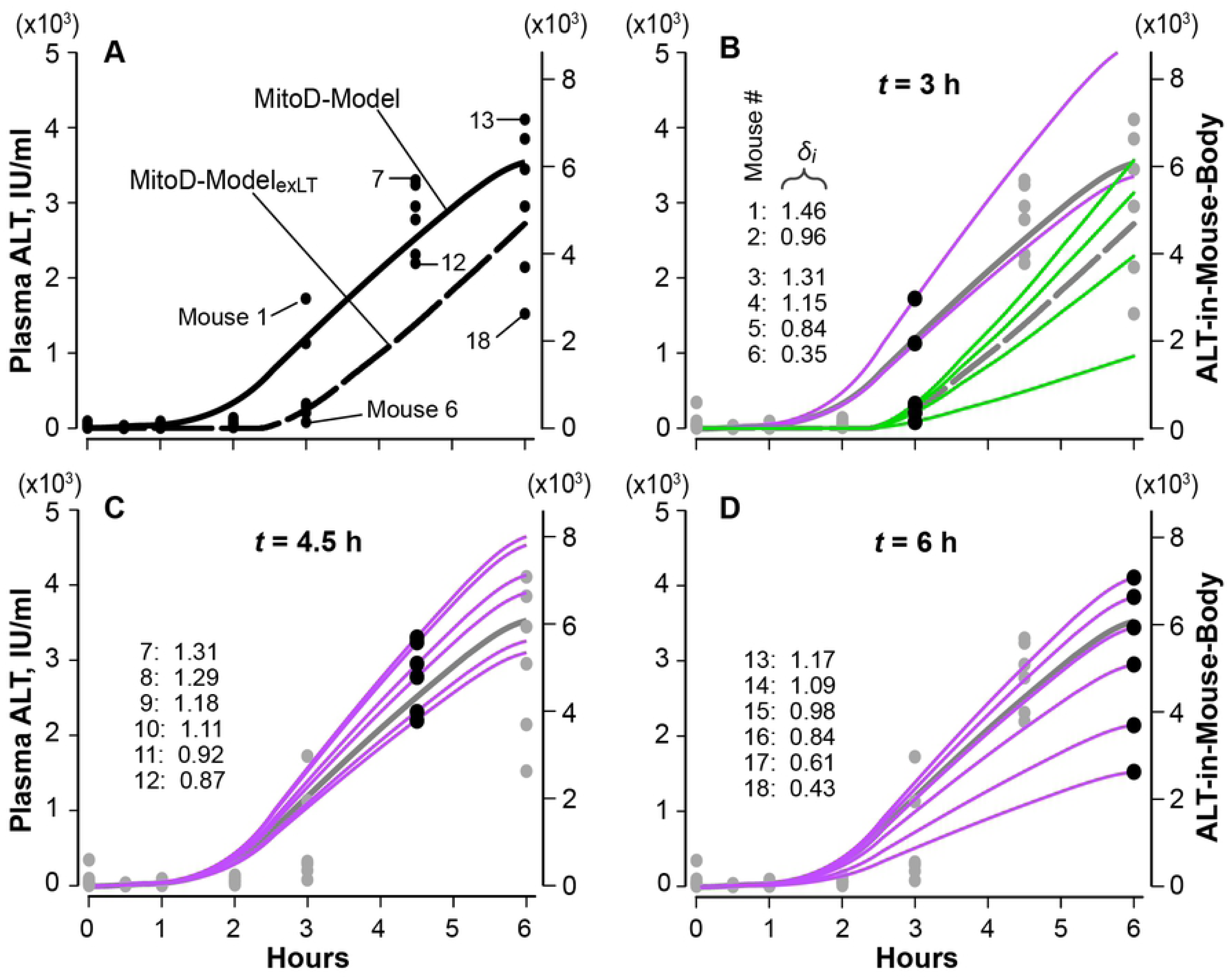
Results from MitoD-Model experiments are scaled to match plasma ALT measures from 18 individual mice. (**A**) Plasma ALT measures in six mice are plotted at the times indicated. We targeted the 18 mice at 3, 4.5, and 6 h; they are assigned numbers in sequence as shown. MitoD-Model: Amounts of ALT-in-Mouse-Body (right axis) are scaled to plasma ALT (left axis) using Equation 1 with S = 1.72 (IU ml-1 ALT-objects-1). MitoD-Model_exLT_ (exLT = extended Lag-Time): The [Min, Max) distribution values that were sampled to determine ALT Release Lag-Time (Fig. 1G) and Necrosis interval for each vHPC are increased by 1600 s (26.67 min). Otherwise, all configuration values are identical to those for MitoD-Model. (**B**) The 3 h validation target values are black; the other values are gray. The solid (dashed) gray ALT-in-Mouse-Body profile is from MitoD-Model (MitoD-Model_exLT_). Individual profiles that match the values for mice 1-6 are obtained by amplifying or diminishing MitoD-Model or MitoD-Model_exLT_ using Equation 2. The *δ_i_* value for each mouse is listed adjacent to the left axis. (**C** and **D**) Designations are the same as in **B**. ALT-in-Mouse-Body values were amplified or diminished to match each of the six individual plasma ALT measures. Statistical measures for the 12 MitoD-Model trials prior to scaling are provided in Supporting S2 Table.

We tested the hypothesis that only shifting the [Min, Max) used by the Leakage Lag-Time distribution to larger values, without changing other parameterizations, would enable the four smallest 3 h plasma ALT measures to also achieve the Similarity Criterion. Results support the hypothesis. We do not know whether the trigger causing the clustering among the 3 h mice also influenced the timing of necrosis-related histopathology. For consistency, because both have common causes, we specified that the [Min, Max) distribution ranges for both Leakage Delay and Death Delay be extended the same. The resulting MitoD-Model variant is designated MitoD-Model_exLT_. We explored several distribution extensions using *S* = 1.72. We achieved the Eq. 2 Individualized Similarity Criterion using a Leakage Lag-Time distribution [Min, Max) = [4300, 19600) ([1.19, 5.44) h; median = 2.875 h) and Necrosis distribution [Min, Max) = [8800, 23200) ([2.44, 6.44) h). Combining the four *δ_i_* for MitoD-Model_exLT_ with the 14 *δ_i_* for MitoD-Model, we obtained *x̄* = 0.992, *s^2^* = 0.0922, *s* = 0.3036, and coefficient of variation = 0.3037.

### Dose-response comparisons: ALT-in-Mouse-Body and plasma ALT measures

McGill et al. [8] also reported early plasma ALT measures following APAP doses of 15, 75, 150 and 600 mg/kg. Does MitoD-Model, parameterized as in Fig. 5, produce reasonably similar ALT-in-Mouse-Body values when dosed with comparable larger and smaller Doses? No.

Measures for Necrosis-Triggered and Necrotic events from Dose-response experiments using MitoD-Model are provided in Fig. 6, A and B. For the “low” and “high” Doses, which correspond to the 150 and 600 mg/kg APAP doses, respectively, the relative temporal patterns within the three Lobular bands are essentially the same as those in Figs. 3 and 4. Focusing on the result at 4.5 and 12 h post-Dose, scaled ALT-in-Mouse-Body values considerably overestimated the corresponding mean plasma ALT measures (Fig. 6, C and D). MitoD-Model is clearly an inadequate explanatory MM for results of both low and high dose experiments. However, while keeping the parameterization of parent MM features (Fig. 1, A-F) fixed, we have a variety of options to improve similarities. From earlier Iterative Refinement Protocol cycles, we knew that the temporal patterns of ALT-in-Mouse-Body values are sensitive to changes in the ALT Leakage Threshold, the Leakage Lag-Time, and Death Delay, so that is where we focused.

**Fig. 6.**
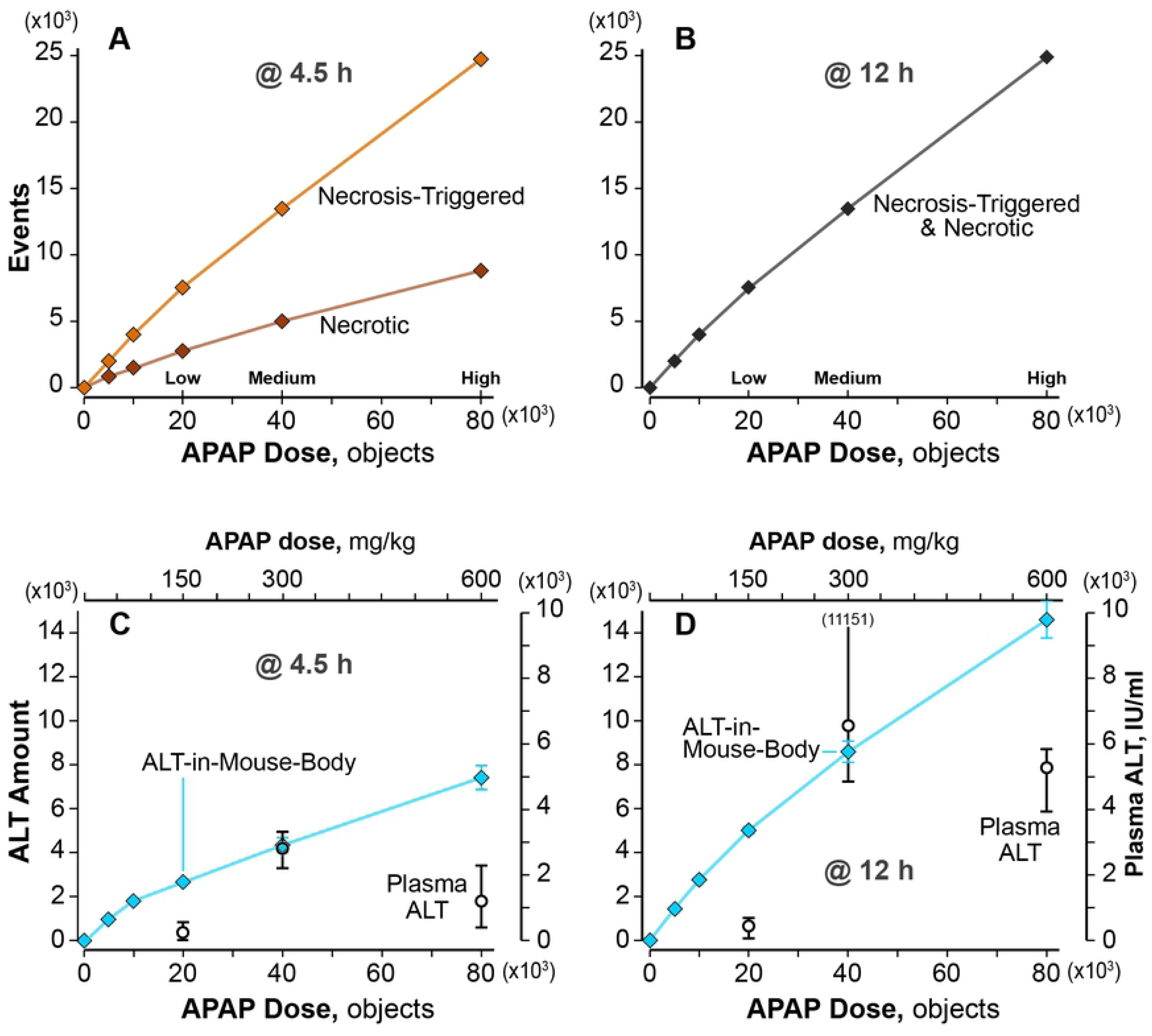
Dose-response relationships for MitoD-Model. (**A**) Average measurements at 4.5 h post-Dose. The three Doses designated low, medium, and high have wet-lab counterparts. (**B**) Average measurements at 12 h post-Dose (results are the same at 12 and 24 h post-Dose). (**C**) Blue diamonds: ALT-in-Mouse-Body values at 4.5 h post-Dose. The corresponding upper and lower whiskers for the two largest Doses span the ALT-in-Mouse-Body range for 12 MC trials. At lower Doses, the symbol eclipses the whiskers. Open circles: mean plasma ALT measures (right axis) for three doses (upper axis) from McGill et al. [8]. The upper and lower whiskers span the plasma ALT range for the six mice. We map the ALT-in-Mouse-Body values to plasma ALT measures using Equation 1 with S = 1.72 (IU ml-1 ALT-objects-1). (**D**) Results at 12 h post-Dose are presented and scaled the same as in C. One of the six plasma ALT measures from the 300 mg/kg experiment is off-scale; the other five values span the range 4,811-6,768 IU/ml.

For the low-Dose, we considered several speculative scenarios. For example, within each vHPC, we could make the value of the ALT Leakage Threshold depend inversely on peak Damage Products (Fig. 3, G and H). There is a huge variety of such seemingly plausible scenarios, but insufficient evidence to shrink the set. To gage how effective such a change might be, we determined the consequences of simply executing MitoD-Model variants in which ALT Leakage Threshold values were increased and decreased. Increasing the Threshold value from 5 to 10 reduced non-Necrotic ALT Release by 88% (90%) at 4.5 h (12 h) and lowered ALT-in-Mouse-Body values. However, the reductions were inadequate. Scaled ALT-in-Mouse-Body values still overestimated the corresponding mean plasma ALT measures because the duration of the Necrosis was unchanged. Necrotic ALT Release—although small—still occurred and that source of ALT significantly buffered the reduction in total ALT that could be achieved by increasing the Leakage Threshold from 5 to 10. Various combinations of increased ALT Leakage Threshold, increased Leakage Lag-Times, and increased Death Delays can be equally effective in improving similarities. However, absent evidence-based constraints and new validation targets, having a large variety of equally plausible MM variants provides no further improvement of explanatory insight.

We can improve similarities at the high Dose using a combination of increased Leakage Lag-Times and increased Death Delays, but here too, having a large variety of equally plausible MM variants provides no further improvement of explanatory insight. Important new knowledge will emerge from discovering plausible MMs capable of explaining plasma ALT measures equally well following all three doses (along with already achieved validation targets). Such MMs will necessarily be significantly more complicated than MitoD-Model. The constellation of those MMs is enlarged dramatically, relative MitoD-Model, and cannot be reduced systematically absent additional wet-lab evidence to serve as new validation targets [19, 20].

## Discussion

The MMs used in generating the results are necessarily complicated. Model mechanism details during execution are entangled within and among levels of virtual Mouse and vLiver organization (Fig. 1). To support clarity in this discussion, we cluster the properties, features, and characteristics of mice (real and virtual) during experiments (real and virtual) into four groups: components, their organization and arrangements (**I**); events, their prerequisites, influence, frequency, order, and sequence (**II**); measurements and measures (**III**); attributes and generated phenomena (**IV**). For the MMs, we have complete, detailed knowledge of **I**. The measures of **IV** can be as rich as needed. Execution produces an observable MM that meets the rigorous definition provided in *Model mechanism requirements*. When measures of **IV** fail to meet prespecified qualitative and quantitative Similarity Criteria, we can discover and explain where, when, and why the failure occurred. Upon achieving the Similarity Criteria, the simulation stands as a challengeable yet tested MM-based theory about abstract, plausible mechanisms (**I**–**IV**) that may have occurred within individuals during the wet-lab experiments. By incrementally strengthening the analogies within **I**, **III**, and **IV**, we provide new knowledge and strengthen the analogies within **II**.

We draw several inferences from the results. Increased plasma ALT measures, following a toxic APAP dose in the targeted mouse experiments are explained best by MitoD-Model, in which ALT Release is driven by both Non-Necrotic Damage and Necrosis, but not by Necrosis alone. The relative temporal contribution of Necrosis and Non-Necrotic Damage during execution (**II**) to ALT-in-Mouse-Body measures (**IV**), which scales directly to plasma ALT measures, changes as simulation time progresses. From a biomarker perspective, it is instructive to note that their relative temporal contributions cannot be inferred from ALT-in-Mouse-Body measures alone.

To match four of the 18 individually targeted plasma ALT measures in Fig. 5, we extended their ALT Leakage Lag-Times and Death Delay intervals (Fig. 5B), while keeping their APAP-induced Hepatotoxicity details unchanged. We infer that the in vivo counterparts of Leakage Lag-Time and Death Delay are sensitive to unidentified non-genetic influences, such as environmental stressors (e.g., within-cage conflict). Such influences limit the reliability of ALT as a biomarker of drug-induced liver injury and likely contribute to interindividual variability. They may also help explain elevated plasma ALT measures absent toxicity.

We limited explorations to extensions of the parent No-ALT-Model (Fig, 1A-1F and Fig. 3) that were suggested by hypothetical mechanistic scenarios described in the cited literature. By doing so, we constricted to a manageable size the constellation of plausible MMs that merited exploration while adhering to the strong parsimony guideline. It is noteworthy that we achieved the stringent validation targets in Fig. 5 without requiring any changes to the No-ALT-Model. Doing so corroborates our claim that the actual spatiotemporal mechanisms causing APAP-induced hepatic injuries within mice following a toxic APAP dose and the virtual counterparts during execution are strongly analogous dynamically. Given the results presented, we extend that claim: at comparable levels of mechanism granularity, the actual spatiotemporal mechanisms causing APAP-induced hepatic injuries in the test mice and giving rise to the targeted plasma ALT measures were strongly analogous to mechanism counterparts within MitoD-Model during execution.

The explanatory power of MitoD-Model following the medium APAP Dose (scales to 300 mg/kg in mice) is significantly eroded for Doses corresponding to 150 and 600 mg/kg of APAP (Fig. 6), indicating that the unfolding and entanglement of crucial temporal features of the mechanism (**II**) in mice is predicated on APAP dose.

We detailed general weaknesses, limitations, strengths, and benefits of both the approach and methods in previous reports [9, 21, 22]. Reliance on analogical arguments and reasoning is both a limitation and strength. Bartha [23] summarizes recent advances in the use of analogical arguments in science and provides guidelines for assessing their strengths and limitations. Because of in vivo measurement limitations, a lack of detailed mechanism-based knowledge, and the fog of multi-source uncertainties, reliance on analogical reasoning and arguments is necessary to make progress in achieving a key objective, which is to begin resolving cause-effect linkages between APAP disposition and simultaneous measurements of ALT in plasma. Because of uncertainties and knowledge gaps, there is still a significant constellation of MMs having similar granularities that meet Requirements (see *Model mechanism requirements*) and are capable of providing equally plausible quantitative explanations of APAP-induced plasma ALT measures in mice.

Consequently, establishing a reliable reverse mapping from ALT measures to particular mechanism features (**I** and **II**) is not yet feasible, and that reality limits the ability of ALT to serve as a mechanism-grounded biomarker of drug-induced liver injury. To improve that reality, we need to shrink the constellation of equally plausible MM-based explanations. We can accomplish that by expanding the set of validation targets to be included within future studies: add temporal measures of a small panel of other putative biomarkers [24], such as mitochondrial RNA-122, glutamate dehydrogenase, nuclear DNA fragments, APAP-protein adducts, and microRNAs. Doing so is also expected to diminish significantly uncertainties currently ascribed to individual variability.

The MitoD-Model is a work-in-progress. It is not intended to become a finished product. For the time being, the MitoD-Model provides a plausible biomimetic model mechanism-based explanation for how the targeted plasma ALT measures are generated. As such, new wet-lab experiments designed to challenge it—falsify it—will yield useful new knowledge, no matter the outcome. If the experiment falsifies a targeted feature, one can use the results from that experiment as new validation targets and follow the Iterative Refinement Protocol to discover a set of MM improvements that enables achieving the new validation targets (while still achieving core validation targets), thus producing a more explanatory, more credible MM. If the experiment fails to falsify the targeted feature, the result strengthens the MitoD-Model analogy. There may be no reason to consider such experiments absent the MitoD-Model and the evidence supporting it. There are multiple features (among **I** and **II**) that may merit a falsification challenge. The following are three examples.

Although the magnitude of each scaling in Fig. 5 differs, the temporal patterns are the same. Most of the events (**II**) driving ALT externalization are completed within the first 3 h (Fig. 3). Consequently, we expect that having sequential measures of plasma ALT from each mouse within 3-6 h post-dose will be most effective in challenging those features. The results will also aid in resolving uncertainties grounded in interindividual variability. Tightly coupling wet-lab and virtual experiments is scientifically sound and is an economical way to concurrently expand explanatory knowledge while chipping away at those uncertainties [7, 9, 21].

The amount of ALT within each vHPC that can be externalized can be another target for falsification. Absent evidence to the contrary, we specified that the amount is the same, independent of PV-to-CV location. Measures of lobular location-dependent ALT activities or ALT gene expression levels (**I**) may (or not) directly challenge that working hypothesis. There is indirect evidence from experiments in rats that does challenge the hypothesis. Gascon-Barré et al. [25] showed that ALT activities in PP hepatocytes are three-fold higher than those in PC hepatocytes. To our knowledge, there are no reports of comparable measurements in mice.

When we Dosed the MitoD-Model with virtual counterparts to 150 mg/kg and 600 mg/kg APAP doses (Fig. 6), the scaled ALT-in-Mouse-Body values failed to mimic the mean plasma ALT measures. Those failures show that the explanatory MMs details must be different following the three doses. Results of exploratory low Dose experiments suggested that using a larger Leakage Threshold and an extended Death Delay range may be sufficient to mimic the referent plasma ALT measures. Whereas, at the high Dose, the results of the exploratory experiments suggested that combining a small Leakage Threshold with extended Leakage Lag-Time and Death Delay ranges may be sufficient. Retaining the same components and features (**I**), one can posit several scenarios for similar feature changes that may enable a single MM to achieve similarities at all three APAP doses. The following are three examples. 1) Make the parameterization of Leakage Threshold, Leakage Lag-Time, and Death Delay ranges (**II**) within each vHPC dependent on the amounts of its Damage Products during the previous simulation cycle. Leakage Threshold would be inversely related to Damage Product amounts, whereas Leakage Lag-Time and Death Delay would increase as Damage Products increase. 2) Specify independent PP-to-PC-dependent values for each of those three features (**I**), analogous to gradients used by the No-ALT-Model, as illustrated in Fig. 1F. Hypothesizing new PV-to-CV location-dependent features is supported indirectly by the fact that over 50% of expressed liver genes exhibit PP-to-PC expression differences [26]. 3) Specify that the Leakage threshold, Leakage Lag time, and Death Delay within a particular vHPC is influenced, via Cell-Cell communication, by the damage state of its neighbors, analogous to the communication features of the model mechanism used by Kennedy et al. [19]. However, we will need new information to select one scenario for further exploration.

For each of the above scenarios, the constellation of plausible yet meaningfully different biomimetic parameterizations will be significant. To improve MM-based explanatory clarity, we need finer-grain measures of release processes to serve as validation targets. With additional evidence-based constraints, in conjunction with new validation targets and strong Similarity Criteria, it is straightforward to use the Iterative Refinement Protocol to alter and add MM features and then select among competing MM-based theories. In so doing, we shrink the plausible constellation of MMs to a manageable size [7, 10, 19].

## Materials and Methods

### Experimental design

The objective of this work is to posit plausible cause-effect linkages between APAP disposition and metabolism and concurrent measurements of ALT in plasma. Until sublobular hepatocyte damage and ALT release can be measured concurrently at multiple times within the same subject, it will be infeasible to establish those linkages in vivo. The alternative approach developed herein is to use virtual experiments to challenge and falsify (or not) many potentially explanatory, biomimetic concretized MM-based hypotheses [22] (see *Model mechanism requirements*). A virtual experiment is a test (a trial) of an extant MM. The approach is based on analogical reasoning [23]. Execution produces an observable mechanism. A consequence of execution that meets the requirements described below is emergence of phenomena that are similar (or not) to prespecified target phenomena, such as pericentral necrosis and ALT externalization. Execution produces a simulation with features that we can measure; those measurements enable testing the hypothesis. If similarities between measures of virtual and real phenomena meet a prespecified Similarity Criterion, then the MM during execution stands as a challengeable yet tested MM-based theory about abstract, plausible mechanism events that may have occurred during the wet-lab experiments.

We experiment on virtual Mice. Their concrete components are strongly analogous to mouse counterparts within and across multiple levels, but only to the extent needed to achieve use objectives and Similarity Criteria [21]. We utilize the virtual experiment protocol outlined by Kirschner et al. [9], as recently enhanced [7, 10]. Mice utilize a previously validated spatiotemporal MM—the parent No-ALT-Model— that explains key early features of APAP hepatotoxicity [23]. Many of the methods used are identical to those detailed by Smith et al. [7, 27]. Nevertheless, to facilitate reproducibility and support descriptions of method extensions and explanations of results, we provide abridged descriptions of methods reused herein.

Because of multisource uncertainties, to achieve the objective it is essential to conduct many narrowly focused virtual experiments to incrementally and systematically shrink the constellation of plausible MM parameterizations capable of achieving validation targets and Similarity Criteria. To do so, we follow an Iterative Refinement Protocol [20, 28, 29], which is a scientific method for falsifying, refining, and validating explanatory MMs.

The Iterative Refinement Protocol begins with a MM, such as the No-ALT-Model, which has achieved several validation targets. We then specify an enhanced Similarity Criterion, or a new feature or phenomenon (e.g., accumulation of ALT-in-Mouse-Body). The immediate goal is that, when scaled, results of executions mimic the validation target sufficiently to satisfy a Similarity Criterion (e.g., the time course of ALT-in-Mouse Body must be sigmoidal). A Similarity Criterion specifies when and how validation target or a Targeted Attribute has been achieved. It can range from qualitative (e.g. event X occurs before event Y; temporal profiles have a sigmoidal shape) to quantitative (e.g. scaled ALT-in-Mouse Body measurements are within 10% of the corresponding plasma ALT measures). MM changes are made that are expected to enable achieving the validation target. The software is reverified. We then specify the configuration and parameterization, and conduct a virtual experiments to test a hypothesis such as this: the time course of ALT-in-Mouse Body will be sigmoidal. Typically, the results of experiments using the initial parameterizations fail to support the hypothesis. That MM is falsified because it cannot be scaled to achieve a given validation target and Similarity Criterion, either because the target phenomenon cannot be generated or we fail to find a parameter set that also satisfies all new and pre-existing Similarity Criteria. Such a falsification completes one cycle of the Iterative Refinement Protocol. The focus then turns to parameterization refinement with the expectation that during the next cycle the validation target will be achieved.

The iterative process of falsification-refinement-validation ensures that the MM is increasingly biomimetic yet parsimonious. The process provides explanatory insight into how, where, and when various referent features may be generated. It also means that MMs are perpetual works in progress, not finished products. We envision MMs being improved incrementally through multiple future rounds of MM challenge and validation against an expanding set of referent measures.

### Making virtual measurements analogous to wet-lab measurements

To strengthen the virtual-to-wet-lab experiment analogy, virtual features and phenomena are measured analogous to how corresponding wet-lab measurements are (or might be) made. At the start of a single execution, the values of multiple parameters and features are Monte Carlo-sampled from prespecified ranges. Most events are stochastic, and probabilities governing them are stated in the specifications. As a consequence of Monte Carlo and probabilistic specifications, measurements can exhibit considerable variability. That variability is intentional to help account for the variability and uncertainty that characterizes wet-lab measurements. Consequently, measurements are averaged across several executions. For this work, the number of Monte Carlo-sampled executions per experiment ranges from 12 (most experiments) to 72 (when small APAP doses were used; the smaller the dose, the greater the measurement variance between executions). The upper and lower whiskers for the ALT-in-Mouse-Body range for 12 MC trials for two APAP Doses are provided in Fig. 6C.

### Model mechanism requirements

We define a mechanism, virtual or real, as entities and activities organized and orchestrated in such a way that they are responsible for the phenomenon to be explained [30]. The MMs studied herein emanate from five demanding requirements that guide software engineering, MM instantiation, and simulation refinements.

1. Five primary characteristics of a biological mechanism – During execution, a MM must exhibit the characteristics of a biological mechanism [31]. 1) For the duration of a virtual experiment, the MM is responsible for a virtual phenomenon that mimics the biological phenomenon to be explained. 2) It has components (modules, entities, etc.) and activities that are 3) arranged spatially and exhibit structure, localization, orientation, connectivity, and compartmentation that are (based on the preponderance of the available evidence) dynamically analogous to biological counterparts. 4) Activities during execution have temporal aspects, such as rate, order, duration, and frequency. 5) The MM has a context, which can include being in a series and/or a hierarchy.
2. Biomimicry – Components and activities are biomimetic [32, 33] according to pre-specified criteria, and they facilitate analogical reasoning [23, 34].
3. Strong parsimony guideline – When scaled, measurements of selected features match or mimic (are strongly analogous to) prespecified Targets (such as the individual plasma ALT measures in Fig. 5) to the extent needed to achieve face validation and specific Similarity Criteria. Adhering to this guideline helps manage the number of equally plausible MMs, while enabling one to increase complexity incrementally. It also facilitates distinguishing a cause from an effect.
4. Emergence – Phenomena measured at a higher level of organization (Fig. 5) arise mostly from local component interactions and phenomena entanglement at a lower level of organization (Figs. 3 and 4).
5. Mobile objects – Each mobile object, such as APAP and ALT, maps to a small amount of their chemical counterparts. Quasi-autonomous components (i.e., software agents such as a SS and a vHPC) recognize APAP and ALT and adjust responses appropriately. For example, a vHPC recognizes that an adjacent APAP has the property *membraneCrossing* = true, and allows it to enter stochastically.

### Mouse components and their organization

The results make the case that, at corresponding degrees of granularity, the MM during execution and the actual APAP-induced hepatotoxicity mechanism may be strongly analogous dynamically within and across multiple lobular levels. That analogy is only as strong as the weakest link in the dynamic networking of MM events during an execution, which depends first on use case-dependent virtual Mouse-to-actual mouse mapping similarities. A Mouse is illustrated in Fig. 1. It comprises Mouse Body, vLiver, and a space to contain Dose for simulating intraperitoneal dosing. It is engineered to facilitate independent replication of experiment results. A vLiver is the number of Monte Carlo-sampled vLobule variants per experiment. vLiver is engineered to be scientifically useful in a variety of usage contexts, but is not intended to (and does not) precisely model a mammalian liver. Rather, it is quantitatively and qualitatively biomimetic during execution in particular ways. It is strongly analogous to actual livers across several anatomical, lobular, and cell biological characteristics. The vLiver in Fig. 1 (and predecessors) achieved qualitative and quantitative validation targets for several different compounds [7, 18, 35, 36]. Consequently, vLiver structure and composition is relatively stable and robust.

Each wet-lab measurement that we seek to mimic, such as APAP hepatic extraction ratio, clearance, metabolite ratios, an intralobular area of necrosis, becomes a Target Attribute. Each Target Attribute is assigned a prespecified Similarity Criterion. A Similarity Criterion specifies the requisite degree of similarity that must be achieved. An example is that, after scaling, the mean measured value of a virtual phenomenon falls within ± 1 standard deviation of the mean target wet-lab value. MM credibility increases by increasing the strength and variety of validation targets achieved.

One vLobule maps to a small random sample of lobular flow paths within a whole liver along with the total volume of associated tissue. It reflects the fact that in vivo APAP and other compounds entering portal vein tracts from blood get exposed to many more hepatocytes than does APAP in blood nearing the central vein. A vLobule comprises a directed graph with a particular SS object at each graph node. Flow follows the directed graph. Graph nodes are organized into three Layers, which map to conventional hepatic zones. Layer 1 = PP zone; Layer 2 = Mid-Zonal; and Layer 3 = PC zone. There are 45 nodes in Layer 1, 20 in Layer 2, and 3 in Layer 3. That structure maps directly to the quasi-polyhedral nature of hepatic lobules. At the start of each execution, all SS dimensions are Monte Carlo-sampled within constraints that enable simulating the wide variety of PP to PC flow paths that were needed to enable the same vLiver to achieve previously described pharmacokinetic Target Attributes for several different drugs [18, 36].

A virtual experiment is a fixed number of Monte Carlo-sampled Mouse executions with a different pseudo-random number seed between each execution. For a 12-Monte-Carlo-execution experiment, the minimum, median, and maximum SS lengths were as follows: Layers 1: 4, 8 and 15; Layer 2: 5, 5.5 and 11; Layer 3: 8, 9 and 10; and vLobule-wide min = 17, mean = 22.5 and max = 36. Intra-Layer edges, primarily within Layer 1 (there are none in Layer 3), mimic interconnections among sinusoids. Numbers of intra- and inter-Layer edges are specified, but their node-to-node assignment are Monte Carlo-sampled for each execution.

Events occurring within a particular SS are dynamically analogous to microscopic referent events occurring within portions of sinusoids and adjacent tissue. Each SS comprises Core, BileCanal (not a factor for this work), and four same-size grids: 1) the Blood-Cell Interface (simply Interface hereafter), 2) Endothelial Cell Space, 3) Space of Disse, and 4) Hepatocyte Space. SS dimensions are Monte-Carlo specified within constraints. Endothelial Cell objects occupy 99% of the Endothelial Cell Space. They contain binders, which can bind APAP and other Solutes nonspecifically. vHPCs occupy 90% of the Hepatocyte space. An APAP object maps to a tiny fraction of an actual APAP dose. APAP Doses are ≤ 100,000 objects. Each simulation cycle (discrete time-step), a fraction of APAP in Body is transferred to PV. From there, Compounds enter Core and Interface spaces at the upstream end of all Layer 1 SSs. Extra-Cellular Compounds percolate stochastically through accessible spaces toward the CV influenced by parameter values that control local flow. Compounds that reach the distal end of Core and Interface spaces are transferred along a connecting edge to another SS. Compounds exit Layer 3 SSs into CV, where they get moved to Body. PV-to-CV gradients provide intra-Lobular location information used by each vHPC. During an execution, each simulation cycle scales to approximately 1 second.

Cells are software agents. Entry and exit of Compounds from Cells is mediated by the Cell according to the Compound’s properties. Endothelial Cells contain binders that bind and release APAP (maps to non-specific binding). Binders map to a conflation of all epithelial cell components responsible for non-specific binding of the referent compound. For a typical experiment, the average number of vHPCs per vLobule was 16,165, with Layer 1 = 10,860, Layer 2 = 4,910, and Layer 3 = 720. vHPCs also contain binders. They map to a conflation of all hepatocyte components responsible for non-specific APAP binding plus all metabolic enzymes responsible for APAP metabolism along with the futile cycle in which APAP is deacetylated to *p*-aminophenol followed by rapid reacetylation back to APAP, even though the cycle is viewed as having little importance in APAP-induced hepatotoxicity in humans and mice [37].

### Model mechanisms that may explain APAP-induced liver Injury

The ALT-in-Mouse-Body phenomenon arises from local component interactions and phenomena entanglement at the lower level of organization illustrated by the measurements in Fig. 4, B-F. Those events trace back through the bottom-level cause-effect linkages occurring within each vHPC (Fig. 3). Events that can occur within a vHPC each simulation cycle are illustrated in Fig. 1E and 1F. vHPC capabilities are identical to those used previously [7]. vHPCs contain four types of physiomimetic modules to control material entry, removal, binding and transformations: *EliminationHandler*, *MetabolismHandler*, *BindingHandler*, and *InductionHandler* (not used in this work). The order of events is (pseudo) randomized each simulation cycle.

The probability of an APAP Metabolism event and the probability that the Metabolite is NAPQI increase threefold from PV to CV. All other Metabolites are lumped together and for simplicity are divided equally between G and S, which map to the APAP glucuronide and APAP sulfate metabolites. The MM uses vLobule-dependent values from each of five gradients illustrated in Fig. 1F. Each gradient is implemented explicitly as a function of distance from PV entrance to the vHPC’s position.

Each vHPC has a location-determined GSH Depletion Threshold value (Fig. 1E and 1F). At early times, there is a 90% chance that as soon as a NAPQI is created, it is eliminated and the GSH Depletion Threshold counter is decremented by 1.0, which maps to depleting a fraction of a hepatocyte’s available GSH. The Threshold is breached when the counter value = 0, which maps to effective GSH depletion. A small or zero Threshold value means that a vHPC is sensitive to NAPQI-caused damage. The iterative process resulting in the location-dependent GSH Depletion Threshold values was described in details by Smith et al. [7].

Each cycle following GSH depletion, there is a 50% chance that a NAPQI will be removed and replaced by (n + 1) Damage Products, one of two types. 1) A MitoD object maps to a conflation of all mitochondria-associated damage products. 2) A nonMD object maps to a conflation of all other types of damage products within a hepatocyte, including those identified with endoplasmic reticulum stress and the unfolded protein response [38]. Each time that one NAPQI is removed, (n + 1) Damage Product objects are created. That number is needed to enable downstream events to be more fine grain, including accounting for accumulation of reactive oxygen/nitrogen species. The value of n is a pseudo-random draw from the uniform [3, 6] distribution.

When the MitoD amount > Necrosis Threshold value (which = 4 for this work), Necrosis is Triggered and the vHPC is designated Necrosis-Triggered. That transition is irreversible. When the Necrosis Threshold is breached, there is a Delay before that vHPC transitions to Necrotic. Transition to Necrotic maps to histologically distinguishable necrosis. To our knowledge, there is no current method available to measure a corresponding transition for hepatocytes in vivo. Because necrosis is a process, there is a delay between the molecular level triggering event(s) and a subsequent time when necrosis becomes clearly detectable in stained tissue sections. That internal process-delay maps to Death Delay in Fig. 1G. There is considerable uncertainty about the timing of triggering events and histological confirmation of necrosis. In Smith et al. [7], Death Delay is a pseudo-random draw from a uniform [Min, Max) distribution = [1.2, 12) h. However, in C57Bl/6 mice administered a 300 mg/kg APAP dose, there is little evidence of necrotic cells prior to 2 h post-dose and little evidence of further necrosis occurring after 12 h. To avoid Necrotic events prior to 2 h during this work, we set Death Delay Min = 2 h (7200 simulation cycles). For the parameterizations used, and a Dose that scales to 300 mg/kg, there are no further Necrosis-Triggered after 6 h post-Dose. We therefore set Death Delay Max = 6 h (21600 simulation cycles). Using a Death Delay distribution = [2, 6) h serves as a predicate to constrain the space searched to identify plausible MMs for ALT release.

Hepatocytes in vivo utilize multiple lobule-location-dependent mechanisms to mitigate or reverse various types of damage. We implemented a single mitigation event by repurposing a Metabolism Module. Damage Mitigation in Fig. 1E maps to a conflation of all actual mitigation/recovery processes, including processes involved in restoring homeostasis. During a Damage Mitigation event, a Damage Product (MitoD or nonMD) may be removed and a Removed object is created and added to the vHPC. The function of Removed objects is to track Damage Mitigation events. The PV-to-CV location-specified probabilities in Fig. 1F for MitoD and nonMD Damage Mitigation events were arrived at following several cycles through the Iterative Refinement Protocol.

### Relationships between parameters governing APAP Toxicity and ALT externalization

The MM for ALT externalization (release) has four parameters: 1) the initial amount of ALT per cell, 2) the damage or leakage threshold, 3) the minimum lag-time, and 4) the maximum lag-time. Parameters 3 and 4 were the main focus of the manuscript. The initial amount of ALT per cell was a constant, and, with the leakage threshold, were not varied to achieve validation. However, the interesting relationship is between parameter 1 and 2, but that relationship is mediated by the amount of damage products. If Leakage Threshold had been larger, e.g., 5 rather than 1, we could still arrive at the same MM phenomena, but it would have required increasing the amplification of nonMD & MitoD. But doing so would have provided no new insights while adding computational costs. In addition, if the initial amount of ALT per cell had been larger, we would need to increase leakage threshold and increase nonMD/MitoD amplification. The amount per cell controls the plateau value in the Body. The results indicate that it is the interplay between threshold and damage that controls the rate of rise of the amount of ALT.

### Limits on mappings

We seek a balance between more detailed biomimicry and the computation programmed into virtual Mice and their methods. A SS does not map directly to a portion of a single sinusoid surrounded by hepatic endothelial cells, hepatocytes, etc. Instead, as described in Hunt et al. [33], events occurring within a particular SS are intended to be strongly analogous to referent events thought to occur within portions of sinusoids and adjacent tissue. The mapping from cylindrical 2D Hepatocyte Space to corresponding 3D configurations of hepatocytes is indirect and not intended to be literal. Instead, as a MM component, it is intended to be adequately (defined in advance) analogous. Because a SS does not map directly to a portion of a single sinusoid, a vHPC cannot map 1:1 to a hepatocyte, although there is a strong functional analogy. Rather, a vHPC at a particular Hepatocyte Space grid point maps to a conflation of relevant hepatocyte functionality at a corresponding PV-to-CV location. Nevertheless, events occurring within a particular vHPC do map directly to corresponding events occurring within hepatocytes at a comparable location.

The scaling factors *S* and *δ_i_* in Equation 1 map the profile shapes between ALT-in-Mouse-Body and plasma ALT measures in mice, establishing the behavioral analogy. Having drawn both structural and behavioral analogies, the above-mentioned limitations are mitigated enough to suggest further mouse studies.

### Technical details

Mice were acquired, cared for, and treated as described in the original studies [8].

The Java-based MASON multiagent toolkit serves as the basis for virtual Mice and many of their components. In earlier work, we referred to virtual Mice as Mouse Analogs (Liver Analogs, Hepatocyte Analogs, etc.) to stress the fact that MM entities are intended to be strongly analogous to their biological counterparts. They do not precisely model the biology. Hunt et al. [22] characterize the spectrum of mechanism-oriented model types being used to help explain biological phenomena. We utilize agent-oriented modeling methods and techniques [39], which allow for complex software entities to be biomimetic in multiple ways. All experiments were run using local hardware and virtual machines [40] on Google Compute Engine, running 64-bit Debian 9. Quality assurance and control details, along with practices followed for validation, verification, sensitivity analyses, and uncertainty quantification are as discussed in [7].

To support verification and validation efforts, all experiments include an “inert” Marker Compound as part of Dose. It acts as a virtual internal standard. It behaves analogous to sucrose in vivo and serves as an indicator of vLobule-structure-Disposition interaction. Absent structural and vLiver component changes, Marker behavior during executions is invariant. Use of Marker is explained further in Supporting S1 Text.

A Mouse is treated as a form of data, using both the implicit schema of Java, JavaScript, and R and the explicit schema of its configurations. Mice and configuration files are managed using the Subversion version control tool in two repositories, one private (Assembla) and another public. The R programming language is used to facilitate analysis and plotting experiment measurements. Values for key vHPC specifications and parameterizations are listed in Supplementary Materials, Table S1. The entire toolchain, including the operating system, configurations, and I/O handling is open-source. The data presented herein, along with the code, are available from project websites (https://simtk.org/projects/aili and https://simtk.org/home/isl/), and are available to be licensed as open data.

### Statistical Analysis

No statistical analyses were needed. Statistical measures of ALT in Mouse Body at 3, 4.5, and 6 h post-Dose from each of the four MM variants are provided in Supporting S2 Table.

## Acknowledgments

We thank the authors of the MASON simulation toolkit and also the creators of the SimTK project-hosting platform for facilitating development and distribution of this project.

## Supporting Information Captions

**S1 Text. Use of a Marker Compound as a Lobule-structure-Disposition Interaction Indicator**

The text describes and provides the rationale for use of a Marker Compound as a type of internal standard during virtual experiments and during most stages of model mechanism development, verification, and validation.

**S1 Table. Description and values for important model mechanism features.**

A list of important configurations/parameters for model mechanistic features specifying the activity and event within vHPCs. The main features are divided into membrane transport of vCompounds, APAP binding/metabolism, damage production/amplification, damage mitigation, necrosis, and ALT externalization (the focus of the main text). In addition, the values listed correspond to the information in Fig. 2 of the main text.

**S2 Table. Statistical measures of ALT in Mouse Body for MM variants.**

The values listed are the minimum, maximum, mean, standard deviation, variance, and coefficient of variation of the unscaled amount of ALT in Mouse Body (12 MC trials) for the four MM variants at 3, 4.5, and 6 h post-Dose. The coefficient of variation is consistent over time and MM variants.

